# Models of microbiome evolution incorporating host resource provisioning

**DOI:** 10.1101/2024.08.05.606723

**Authors:** Yao Xiao, Teng Li, Allen Rodrigo

**Affiliations:** School of Biological Sciences, The University of Auckland, Auckland 1010, New Zealand

**Keywords:** Microbiome, Model, Evolution, Resource provisioning

## Abstract

Multicellular hosts and their associated microbial partners (i.e., microbiomes) often interact in mutually beneficial ways. Consequently, hosts may choose to allocate resources to regulate and recruit appropriate microbes. In doing so, hosts may incur an energetic cost, and in turn, these costs can affect host fitness. It remains unclear how hosts have evolved to balance the costs of expending resources to manage their microbiomes against the benefits that might accrue by doing so. We extended a previously-developed agent-based computational model of hostmicrobiome evolution by incorporating a resource provisioning process, whereby hosts provide resources to support beneficial microbes and suppress harmful microbes, with attendant fitness costs. Our results indicate that the ways and sources from which a host acquires microbes are crucial factors influencing the host’s willingness to provide resources, to regulate microbiome composition. The intensity of resource provisioning will depend, in part, on how much of their microbiome hosts contribute to, and obtain from, their environment: when hosts that engage in resource provisioning contribute a high percentage of their microbiome to the environment, then there is less evolutionary imperative for other hosts to also provide resources. Since resource provisioning incurs a fitness cost to the host, over evolutionary time, resource provisioning will not be favored. Additionally, if selection at the microbial level is not sufficiently strong, and the host obtains a high proportion of microbes from the environment, then the higher the proportion of beneficial microbes in the environment, the less the host is willing to provide resources.

## Introduction

Many multicellular organisms host microorganisms that influence hosts’ fitnesses in various ways [1]. Microorganisms can help the host digest food [2], promote the development of the immune system [3], prevent disease [4], facilitate adaptation to the environment [5], provide vital nutrients [6] and even have an impact on the psychological state of the host [7].

Zeng et al. developed neutral [8] and selection-based [9] computational models exploring the evolution of hosts and their associated microorganisms – their microbiomes – over many generations of hosts. The latter study investigated the changes in microbiome diversity and fitness of the host and microbiome under different intensities of host and microbial selection in conjunction with different modes of microbial acquisition (i.e., horizontal and vertical transmission).

In the relationship between host and microbiome, it is not only the microbiome that influences the host – the host also has a strong influence on the microbiome [10]. Hosts often provide favorable environments and resources for beneficial microorganisms; conversely, they may also employ various mechanisms to suppress harmful microorganisms. Legumes provide essential nutrients and energy for rhizobia [11]; corals provide shelter for zooxanthellae and substrates (e.g., *CO*_2_) for photosynthesis [12]; the Hawaiian bobtail squid (*Euprymna scolopes*) has evolved an organ specifically for bioluminescent bacterium (*Vibrio fischeri* ) as a habitat [13]. Carpenter ants can reduce the number of bacterial symbionts through immune system regulation during their development [14].

It is reasonable to assume that hosts that actively regulate their microbiomes sustain an energetic cost in doing so. This cost may be because hosts have to sacrifice some of their resources to construct new environments or to suppress parts of their immune systems, thus risking infection. Over evolutionary time, the resources that hosts sacrifice can be modelled as a fitness cost. The host therefore needs to balance the cost incurred by resource provisioning against the fitness benefits that accrue with the retention of beneficial microbes and/or the suppression of harmful microbes. How do hosts balance these costs and benefits?

In this paper, we extend one of the selection models by Zeng et al. – the trait-mediated microbial selection model (TMS) [9] – to include a host resource provisioning process (RPP) whereby hosts incur fitness costs to provide resources to recruit beneficial microbes and suppress harmful microbes. Our results indicate that the intensity of resource provisioning will depend, in part, on how much of their microbiome hosts contribute to, and obtain from, the environment: when hosts that provide resources to retain beneficial microbes contribute a high percentage of their microbiome to the environment, then there is less evolutionary imperative for other hosts to also provide resources, particularly if these hosts acquire a large percentage of their microbiome from the environment. Since resource provisioning incurs a fitness cost to the host, over evolutionary time, resource provisioning will not be favored. Additionally, if selection at the microbial level is not strong enough, and the host obtains many microbes from the environment, then the higher the proportion of beneficial microbes in the environment, the less the host is willing to provide.

## Model

Zeng et al. [9] developed a forward-time computational agent-based model for simulating the evolution of hosts and microbiomes under selection. This model extends an earlier neutral model also developed by Zeng et al. [8]. For both neutral and selection models, each iteration of the simulation is equivalent to a discrete generation of hosts. The size of the host population remains constant from one generation to the next. In the absence of selection, hosts have equal probabilities of producing offspring in the succeeding generation, whereas under selection, the probability that a host will have offspring is a function of its relative fitness.

At each generation, each host acquires a percentage of microbes from its (single) parent in the previous generation (*x* %) and from the environment (1-*x* %). When *x* = 0, all microbes are obtained from the environment; when *x* = 100, all microbes are obtained from the parents. The model also allows previous generations of hosts to contribute some fraction of their microbiomes to the environment. The environmental microbial community is constituted by the acquisition of *y* % from the collective host microbial community in the previous generation and (1- *y* %) from a fixed environment. In a fixed environment, the frequencies of all microbial species/taxonomic groups (for simplicity, we will refer to these as Operational Taxonomic Units, or OTUs) remain the same in each generation (this may be equivalent to a biological system where there may be continuous input of microbes from an allocthonous source, e.g., a river or estuary, or where there is ongoing regeneration of autocthonous microbes, e.g., decomposing wood and litter in a forest). When *y* = 0, the environment is completely unaffected by the previous generation of hosts. Conversely, when *y* = 100, all microbes in the environment are inherited from the previous generation of hosts. Hosts in the first generation obtain microbes directly from the fixed environment by sampling from a uniform distribution.

In the computational model with selection, each OTU is assigned a phenome composed of a fixed number of traits. Each trait can affect the fitness of OTUs and hosts independently; thus, each trait is assigned two values: one represents the effect of the trait on the fitness of hosts, and the other represents the effect of the trait on the fitness of the OTU. The values that these traits can take are -1, 0, or 1, where -1 indicates that the trait has a negative impact on fitness (of hosts or OTUs), 0 indicates that the trait has no effect on fitness and is therefore neutral, and 1 indicates that the trait has a positive effect on fitness. Based on these traits, the contributions of each OTU to itself and to its host can be calculated.

The frequency of each OTU within the microbiome of an individual host is a function of its frequency when it is first acquired, either from the host’s parent or the environment, and the fitness of its phenome. Host selection (HS) impacts the survival and reproductive ability of the host, and can be calculated based on the composition of the microbiome within the host and the contribution to the host fitness of each OTU. All things being equal, the higher the frequency of beneficial microbes in the host, the higher the probability of reproduction. Zeng et al. [9] considered two types of microbial selection (MS): first, under trait-mediated selection (TMS), a given trait makes the same contribution to microbial fitness, regardless of which host an OTU that possesses that trait is in; in contrast, under host-mediated selection (HMS), the contribution that a given trait makes to an OTU’s microbial fitness is modified depending on which host the OTU is present in. HMS is meant to capture the process whereby hosts have certain heritable attributes that favour (or disfavour) OTUs with some traits.

In this present study, we focus exclusively on TMS, and we have introduced a resource provisioning process (RPP) whereby we assign a specific trait to each host, representing the amount of resources it provides to promoting beneficial microbes and suppressing harmful microbes (Fig. 1). This trait can also be inherited from parent to offspring. At the start of each simulation, the trait is assigned a value, *k*, ranging from 0 to 1 sampled from the uniform distribution. The value *k* represents the extent to which the host is willing to expend resources to shape the microbiome it carries. When *k* = 0, the host is unwilling to allocate any resources to the microbes. When *k* = 1, the host is willing to allocate as much resource as possible to the microbes. Additionally, in this model, hosts can recognize whether microbes are beneficial or non-beneficial and respond differently based on the value of *k*. For beneficial OTUs (contribution to the host fitness *>* 0), the host reallocates its own fitness to the OTU’s microbial fitness. For neutral OTUs (contribution to the host fitness = 0), the host does not contribute any resource, and thus has no impact on the fitness of these OTUs. For non-beneficial OTUs (contribution to the host fitness *<* 0), the host sacrifices its own fitness to decrease microbial fitness.

**Figure 1:**
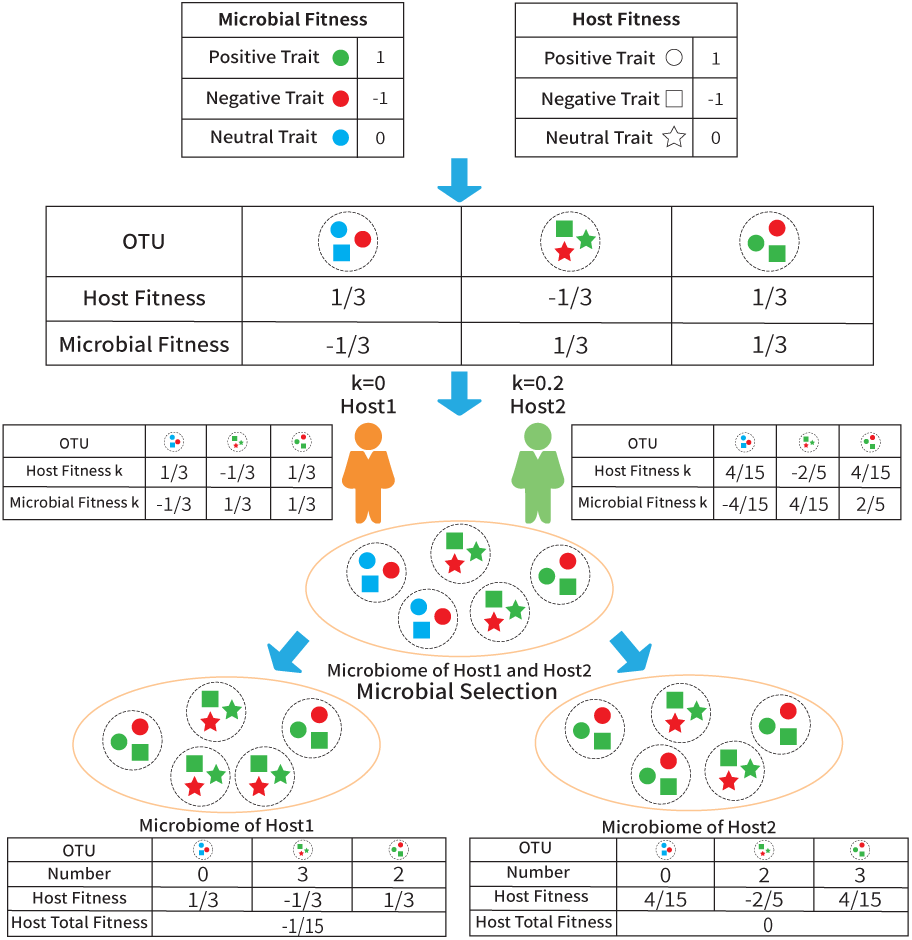
An example of resource provisioning process (RPP). The diagram shows how RPP operates on hosts with different values of *k*. In this example, there are three OTUs, each characterized by three traits. The color of the traits represents their impact on contribution to the microbial fitness (green for positive, red for negative, and blue for neutral). The shape of the traits represents their impact on host fitness (circle for positive, square for negative, and star for neutral). Host *1* does not sacrifice any of its fitness to promote beneficial microbes and suppress non-beneficial microbes within its microbiome (*k* =0). Host *2* allocates a small portion of resources (*k* =0.2) to create a favorable environment for the relevant microbes. After RPP, the contribution to the host fitness of OTUs within Host *1* remains unchanged. However, in Host *2*, the contribution to the host fitness of all OTUs decreases, but the contribution to the microbial fitness of positive microbes increases, while the contribution to the microbial fitness of negative microbes decreases. Subsequently, microbial selection occurs within the hosts, resulting in changes to the microbiome. In Host *1*, the proportion of negative microbes in the microbiome increases while the proportion of positive microbes decreases. On the other hand, in Host *2*, the proportions of negative and positive microbes remain unchanged. After comparing Host *1* and Host *2*, we can see that without the inclusion of RPP, negative microorganisms would have higher fitness compared to positive microorganisms, and the host total fitness would decrease with each generation as the negative microorganisms increase. However, with the incorporation of RPP, this situation improves. As a result, host total fitness of Host *2* is higher than Host *1*.

We conducted two different sets of simulations. In the first set, we set *k* = 0. This is equivalent to a simulation without RPP. We did this as a control so that we could compare our results to those obtained by Zeng et al. [9]. In the second set, we set a uniform random *k* from 0 to 1 to each host at the start of each simulation. These simulations allow us to compare microbiome diversities and host/microbe fitnesses in the presence and absence of RPP.

## Results

### Microbial Diversity

Results of our simulations in the absence of RPP (Fig. 2*a*) recapitulate those of Zeng et al. [9]: under nearly neutral conditions (very low intensity of MS, *S_MS_* = 1.1), a significant decrease in *α*-diversity can be observed only when the contribution of parents to offspring and the environment is high (*≥* 90%). Furthermore, *α*-diversity exhibits a clear decreasing trend with increasing intensity of MS (as we move down the rows of heatmaps in Fig. 2*a*). However, increasing the intensity of HS does not result in significant changes in *α*-diversity.

**Figure 2:**
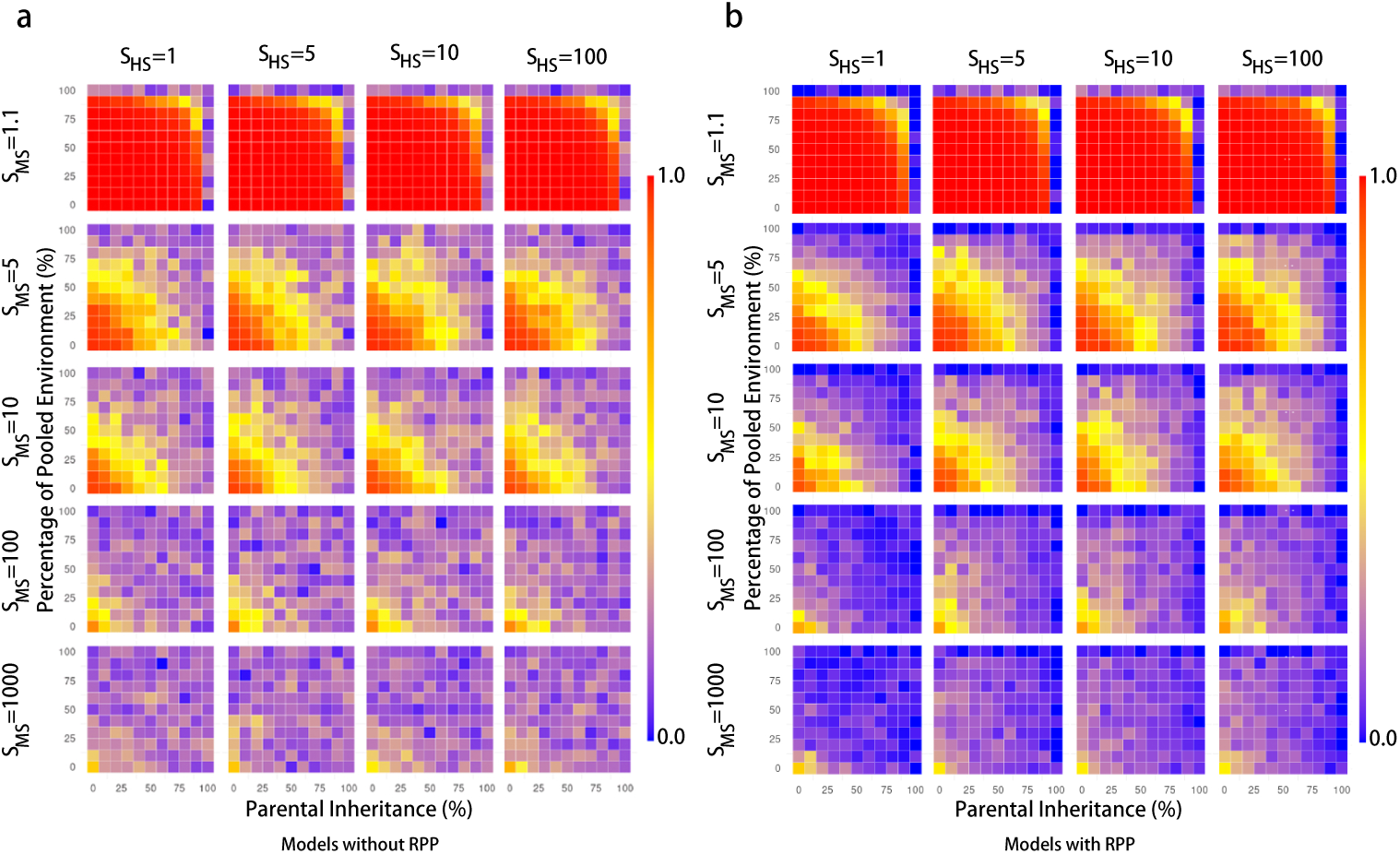
Pattern of *α*-diversity without (*a*) and with (*b*) RPP. Each heat map represents its corresponding combination of HS and MS. There are five levels of MS: 1.1, 5, 10, 100, 1000, and four levels of HS: 1, 5, 10, 100. For each heat map, the horizontal axis is the parental contribution to the host (i.e., parental inheritance, PI), and the vertical axis is the parental contribution to the environment (i.e., the percentage of pooled environment, PE). The scale of the horizontal and vertical axes of each heatmap is linear, from 0 to 100. The color bars on the right side represent the corresponding diversity values (blue for low diversity, yellow for medium diversity, and red for high diversity) This, in turn, leads to a decrease in *α*-diversity.

Although the overall patterns are similar after incorporating RPP, there are slight variations in *α*-diversity (Figure 2*b*): with RPP *α*-diversity decreases even more than when RPP is absent, especially as the proportion of parental contributions to the host and environment increases. As we show below, the inclusion of RPP leads to a decrease in the number of negative microbes,and an increase in the number of beneficial microbes.

Similar to the results from Zeng et al. [9], *γ*-diversity and *α*-diversity exhibit a high degree of concordance (*Supplemental files 1*, Fig. S1), while *β*-diversity remains consistently low under all conditions (*Supplemental files 2*, Fig. S2).

### Resource Provisioning parameter, *k*

By incorporating the RPP parameter, *k*, our simulations show the amount of resources the host contributes to microbes, or equivalently, the amount of fitness the host will sacrifice (Fig. 3). When the microbiome makes no contribution to host selection, (i.e., *S_HS_* = 1), the microbial contribution to host fitness of all traits = 0. Under the RPP model, *k* has no effect on this value, since there is no net loss to host fitness, regardless of the value of *k*. Therefore, *k* remains a uniform random number between 0 and 1, with an expected value of 0.5.

**Figure 3:**
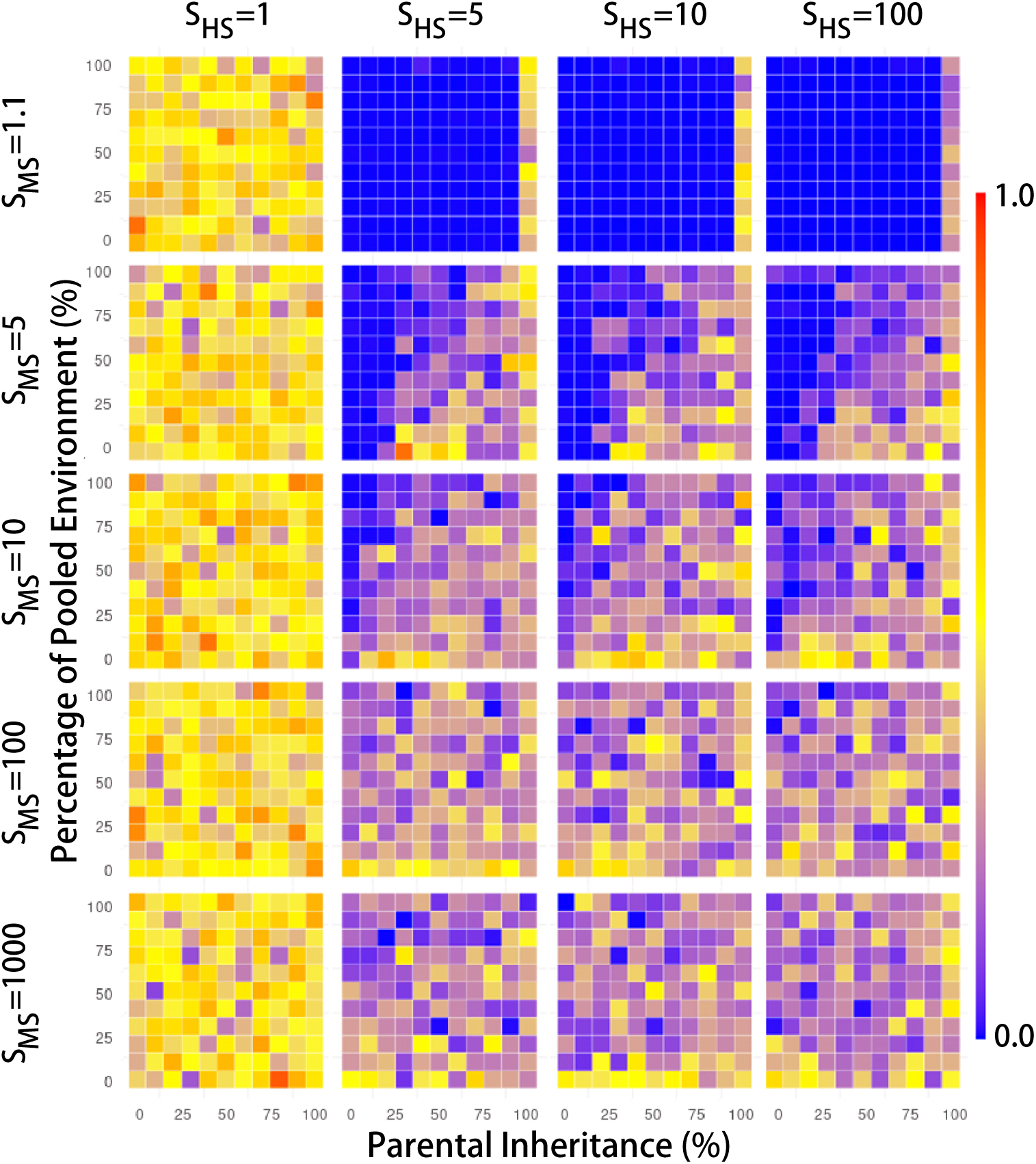
Pattern of *k* in the presence of RPP. This figure is plotted in the same way as Fig. 2, but the color represents the corresponding values of *k*.

When there is almost no MS (*S_MS_* = 1.1; the first row of heatmaps in Fig. 3), *k* is close to 0, regardless of the value of *S_HS_* . This is because even though a value of *k >* 0 implies a loss of host fitness, the investment of hosts in the microbiome has very little effect on the differences in fitness of beneficial, neutral, or non-beneficial microbes. Therefore, hosts that sacrifice fitness (i.e., *k >* 0) do so for very little, or no, increase in reproductive success. Hence, over evolutionary time, selection favors hosts that contribute no resource to their microbiomes. (Note that the exception to this is when parental inheritance is 100%, the last column in each of the heatmaps in the row when *S_MS_* = 1.1. In this case, even though the intensity of MS is weak, the offspring can inherit the entire microbiome from the host, and there is value in retaining beneficial microbes, and removing non-beneficial microbes).

When *S_MS_ >* 1.1 and *S_HS_ >* 1, the situation changes. Although the intensity of HS does not significantly change the distribution of *k*, for any given value of *S_MS_*, as the intensity of MS increases, *k* also increases. Because *k* increases the fitness of microbes that are beneficial to the hosts, and decreases the fitness of microbes that are harmful to the host, increasing *k* leads to an increase in the fitness differences between beneficial and non-beneficial OTUs. In this scenario, the number of beneficial microbes to the host increases rapidly, resulting in higher returns on the investment of hosts in microbes.

Finally, when considering the effect of parental inheritance on *k*, we can see that the value of *k* increases as the percentage of parental inheritance (PI) increases. If the host can pass on the microorganisms obtained through RPP to the offspring, the host is willing to provide more resources, tending to a higher *k* (see, for instance, Fig. 3 lower righthand corners of heatmaps, where values of *k* tend to be higher). On the other hand, *k* tends to decrease as the percentage of hosts’ contribution to the pooled environment (PE) increases and when hosts acquire a substantial portion of their microbiomes from the environment (see upper left-hand corners of heatmaps in Fig. 3). A higher PE indicates that the environment is constituted by a higher percentage of microbes obtained from the previous generation of hosts than from a fixed environmental pool. Since other hosts in the environment recruit substantially from the same environment, they are able to recruit beneficial microbes deposited by hosts that have *k >* 0, without themselves having to sacrifice fitness. The hosts that do not participate in RPP will increase in frequency, and over evolutionary time, selection acts to reduce the frequencies of hosts with high values of *k*.

To confirm this, we performed a simple linear regression analysis for each of these two independent variables (parental inheritance, PI, and parental contribution to the environment, PE) against the dependent variable *k* (Fig. 4*a* and Fig. 4*b*). As expected, when there is no HS (*S_HS_*= 1) or very weak MS (*S_MS_* = 1.1), there is no significant correlation between *k* and these two factors (PI and PE).

**Figure 4:**
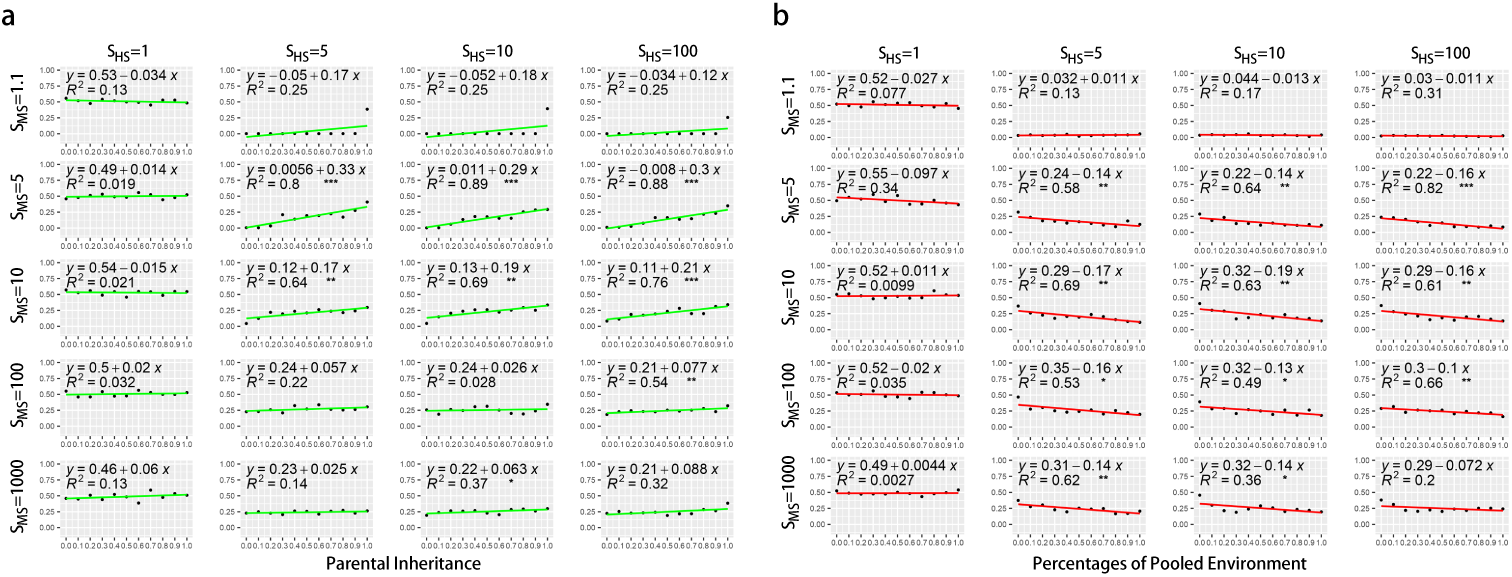
Simple linear regression analysis of *k* s and parental contribution to hosts (a) and environment (b). Each scatter plot represents its corresponding combination of HS and MS. MS has five levels: 1.1, 5, 10, 100, and 1000; HS has four levels: 1, 5, 10, and 100. For each scatter plot, the horizontal axis is the parental contribution to the host, from 0 to 1. The vertical axis is *k*, and each point is obtained by averaging the data of different parental contributions to the environment with the same parental contribution to the host. The red line represents the model obtained by one-dimensional linear regression. The * in the graph represents the p-value (*** represents *p <* 0.001, ** represents *p <* 0.01, * represents *p <* 0.05). Since the purpose of plotting this figure is to visualize the trends of *k*, rather than strict hypothesis testing, we chose not to apply a Bonferroni correction to the p-values.

When *S_MS_ >* 1.1 and *S_HS_ >* 1, there is a tendency for *k* to be positively correlated with PI (this is most obviously seen when MS is moderately intense), confirming that an increase in parental transmission of microbes is associated with an increase in the amount of resource a host is willing to provide to retain beneficial microbes and suppress harmful microbes.

When *S_MS_ >* 1.1 and *S_HS_ >* 1, *k* is negatively correlated with PE (this is also most obviously seen when MS is moderately intense). Again, this confirms that over evolutionary time, hosts that provide resources to beneficial microbes and subsequently deposit these microbes in the environment are at a selective disadvantage. The evolutionary outcome of this is that hosts with lower values *k* will dominate the population.

Although a higher intensity of MS can lead to a relatively higher *k* (i.e., *PI* = 0, 10, 20), this is not always the case. We speculate that factors influencing *k* may extend beyond the parental contribution rate and selection intensity alone. It is possible that the initial composition of the host’s microbiome also exerts an influence on *k*. To explore this further, we conducted an all-subsets regression analysis of these factors, with the dependent variable as *k*, and the five independent variables as the initial proportion of beneficial microorganisms, PI, PE, the intensity of MS, and the intensity of HS. The most influential variable on *k* is the initial proportion of positive microorganisms in hosts, which is negatively associated with *k*, i.e., when the initial proportion of positive microorganisms is high, then *k* tends to be low. The other influential variables, in order of the magnitudes of their contributions to the variation in *k*, are the contribution rate of the parents to the offspring (positively associated with *k* ), the contribution rate of the parents to the environment (negatively associated with *k* ), the intensity of MS (positively associated with *k* ), and the intensity of HS (negatively associated with *k* ) (*p <* 0.05).Thus, if the host begins with a higher proportion of positive microorganisms in its initial microbiome, it does not need to allocate additional resources to its securing the services of beneficial microbes.

### Composition of Microbiomes

How does RPP affect microbiome composition over evolutionary time? To answer this question, we graphed the composition of the microbiome at the end of the simulations (Fig. 5). When MS intensity is very weak (*S_MS_* = 1.1), the composition of the microbiome is highly variable with respect to the distribution of beneficial, neutral, or harmful microbes. As the intensity of MS increases (*S_MS_ >* 1.1), the percentage of positive microorganisms increases. As noted above, the host’s fitness sacrifice increases the microbial fitness of beneficial microbes and reduces the microbial fitness of harmful microbes, creating larger differences in microbial fitness between the two categories of microbes. The high intensity of MS leads to a further expansion of beneficial microorganisms.

**Figure 5:**
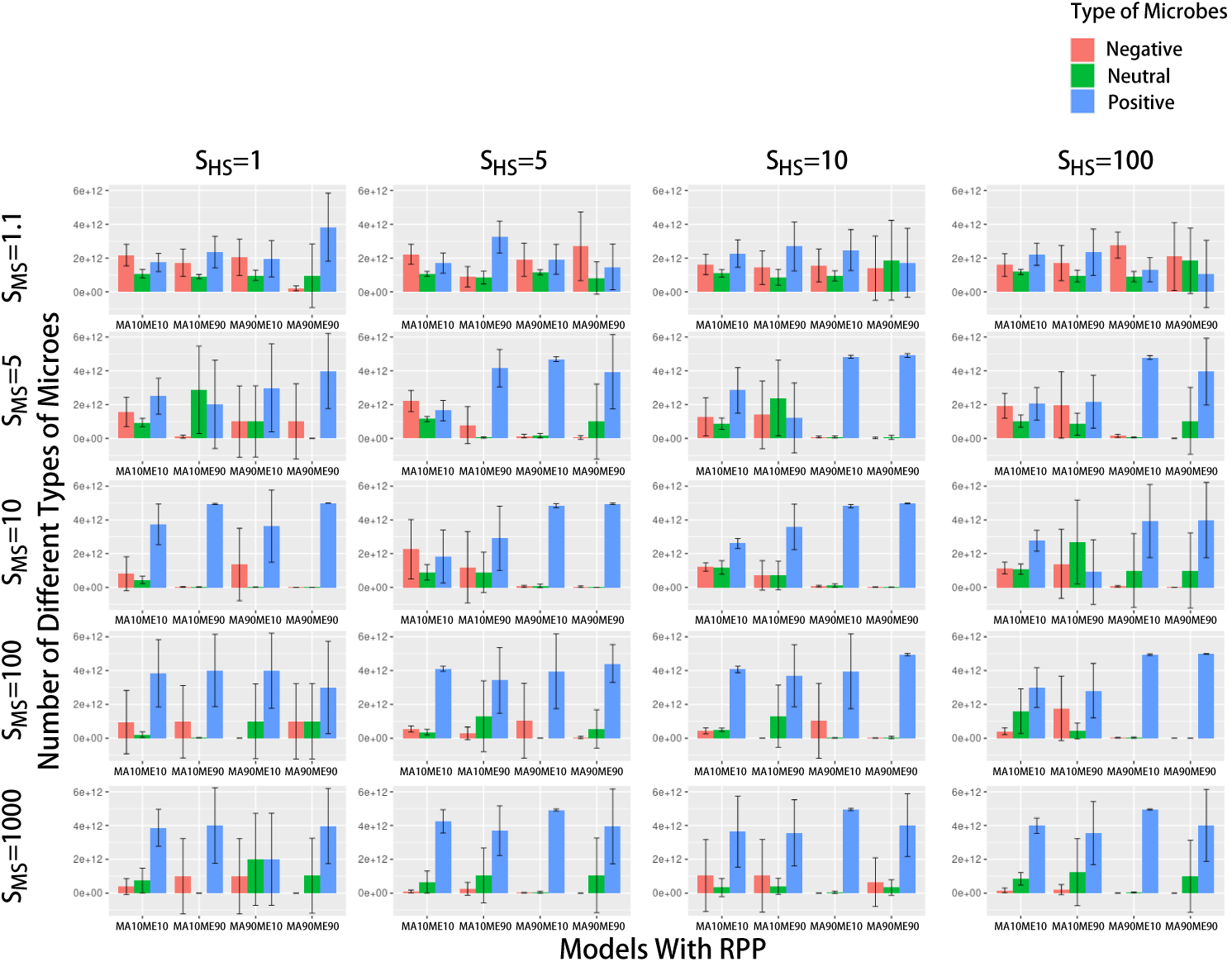
Composition of microbiomes under different models. Each barplot represents its corresponding combination of HS and MS. There are five levels of MS: 1.1, 5, 10, 100, 1000; and four levels of HS: 1, 5, 10, 100. For each barplot, the horizontal axis represents different combinations of parental contribution rates: *MA10ME10*, *MA10ME90*, *MA90ME10*, *MA90ME90*. *MAnMEm* represents a parental contribution of n% to the host and m% to the environment. The vertical axis represents the microbial proportion, and each bar represents the average value of five repetitions. The lines at each bar represent *±* one standard deviation. Red represents microbes that have a harmful effect on the host (negative microbes), green represents neutral microbes, and blue represents microbes that have a beneficial effect on the host (positive microbes).

The percentage of positive microorganisms also increases with an increase of PI because a high PI means that the host can inherit more beneficial microbes from its parent. When PI is low (*MA10* ), the host acquires most of the microorganisms from the environment. Nonetheless, it is worth noting that neutral or even harmful microorganisms do not become extinct in many cases, even though PI is high (*MA90* ).

### Fitness of Hosts

Unsurprisingly, RPP not only increases the proportion of positive microorganisms but also increases the average fitness of the host (Fig. 6*a*). Compared to simulations without RPP, applying HS leads to an increase in hosts’ average fitness. This effect is observed with increasing intensity of HS and PI. We conducted an Analysis of Covariance using the change of host fitness as the dependent variable, the presence/absence or RPP, PI, and the intensity of HS as independent variables. The results confirmed there is a significant increase in host fitness in the presence of RPP (*p <* 0.001). The increase in host fitness is also associated with increasing PI and intensity of HS (*p <* 0.001). In essence, although the host sacrifices fitness as part of RPP, this is compensated by the increase in fitness contributions of beneficial microorganisms. This is especially true when PI is high; in this case, parents leave more beneficial microorganisms for offspring.

**Figure 6:**
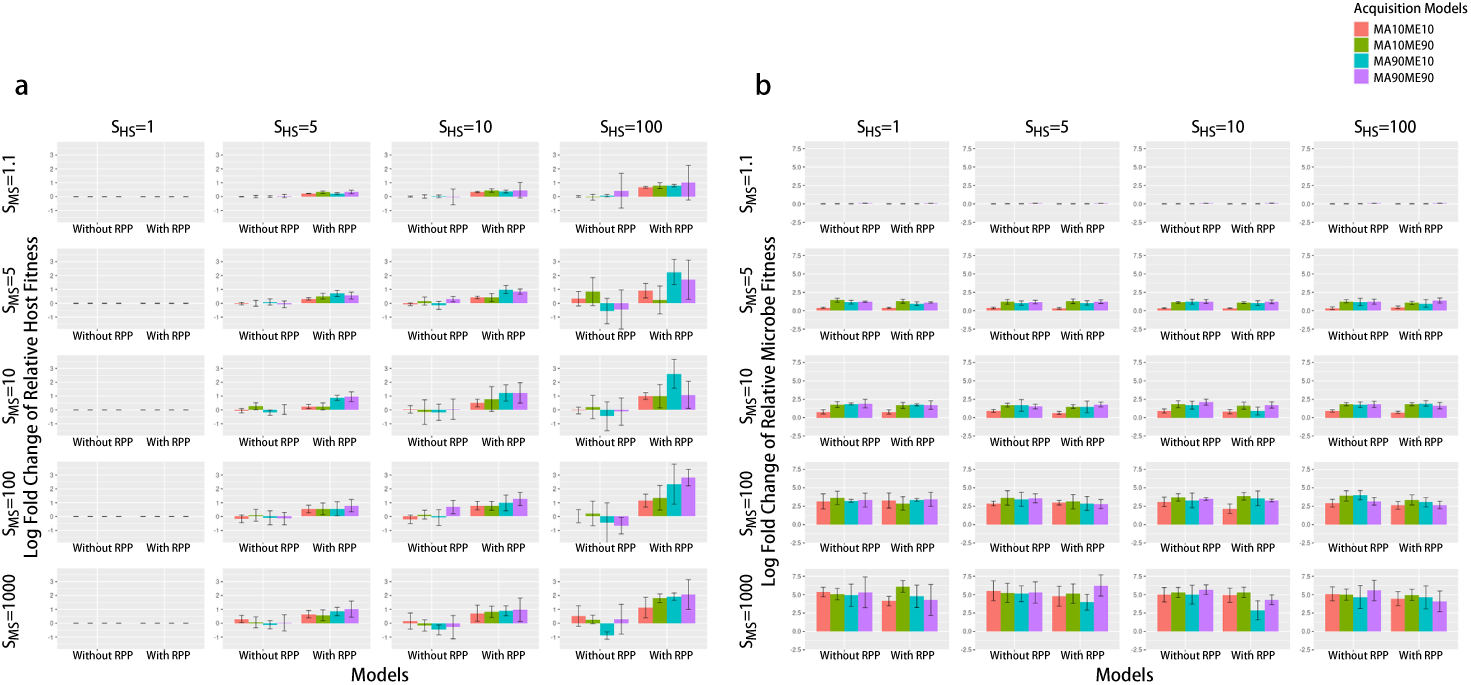
The average log-fold relative changes in host fitness (a) and microbial fitness (b) at the start and end of the simulations with and without RPP. Each barplot represents its corresponding combination of HS and MS. There are five levels of MS: 1.1, 5, 10, 100, 1000; and four levels of HS: 1, 5, 10, 100. The vertical axis represents the logarithmic change in average host fitness relative to the initial level. The lines at each bar represent *±* one standard deviation. The horizontal axis represents whether RPP was included. Different colors represent different combinations of parental contribution rates.

## Discussion

We extended the selection model developed by Zeng et al. [9] to incorporate the process of host resource provisioning. Under resource provisioning, hosts invest resources (at a fitness cost) to create an environment conducive to recruiting beneficial microorganisms and to suppressing harmful microorganisms.

Our results show that the *α*-diversity of the microbiome decreases after the inclusion of the host’s resource provisioning process (RPP). In other words, if the host provides resources to create a specialized environment for the resident microbiome, it can lead to the elimination or failure of colonization of microorganisms that are harmful to the host while promoting the colonization and survival of beneficial microorganisms. As a result, the *α*-diversity of the microbiome is reduced. Indeed, in biological systems, we can find examples of such phenomena. For instance, the human gut provides specific temperature and pH conditions that only allow the colonization and proliferation of certain microorganisms [15, 16]. As a result, the gut microbiome is primarily composed of the Bacteroidetes and Firmicutes bacteria, along with the archaeon *Methanobrevibacter smithii* [17, 18]. Overall, our simulation results suggest that if the host shapes a specialized environment, it can lead to a decrease in the *α*-diversity of its microbiome.

The “willingness” of the host to provide resources in shaping the environment (i.e., explicitly, an increase in the average RPP parameter *k* ) increases with a higher parental contribution to the offspring. However, the value of *k* decreases with a higher parental contribution to the environment, particularly when hosts acquire a large proportion of their microbes from the same environment. In this case, over evolutionary time, hosts are less inclined to provide resources in shaping their microbiomes. This situation is somewhat similar to the “Black Queen Hypothesis” [19]. The similarity lies in the initial stage of our simulation, where hosts with different resource expenditures are present. High-resource-provisioning hosts release a larger number of beneficial microorganisms into the environment, providing an advantage to low or non-resource-provisioning hosts and their descendants. Non-resource-provisioning hosts conserve resources (i.e., they have low or zero value of *k* ) and gain a fitness advantage.

When focusing on the composition of the host microbiome, we find that in the presence of host resource provisioning, the proportion of microorganisms beneficial to hosts increases, as expected. An example of this is seen in the Hawaiian sepiolid squid, *Euprymna scolopes*, which has evolved a specialized structure called the light organ that facilitates the establishment and maintenance of a population of the bioluminescent marine bacterium *Vibrio fischeri* [20, 21]. As a result, the concentration of *Vibrio fischeri* within the light organ is much higher than in the surrounding seawater [22, 23].

Our results also indicate that if the host can acquire the majority of microorganisms from their parents, parents are more likely to invest resources in beneficial microbes, leading to an increase in host fitness. Research has shown that vertical transmission of microorganisms from the parent to offspring is widespread [24]. For example, in sponges, up to 33 microbial clusters spanning ten bacterial phyla and one archaeal phylum are vertically transmitted [25–28]. In the Cynidae family of stinkbugs and Coreidae family of leaf-footed bugs, offspring acquire microorganisms by consuming the feces of their mothers [29, 30]. Female *Sceloporus virgatus* lizards can transfer beneficial microorganisms to their offspring during the egg-laying process to protect the eggs from fungal infections [31].

Our findings in microbiome fitness are also intriguing. Under the condition where the host provides a special environment for microorganisms, the native fitness of the microbiome decreases. Considering the composition of the microbiome, this suggests that microbial fitness is not the sole criterion for microbial survival within the host – under RPP, the host plays an active role in boosting microbial fitness of beneficial microbes that may otherwise be less fit than other microbes. In the human vaginal microbiome, 90% of the microorganisms are *Lactobacillus* [32, 33]. *Lactobacillus* not only adapts to the low pH environment of the human vagina but also produces bacteriocins, which help resist the invasion of pathogens [34].

By necessity, our models make certain simplifying assumptions. Our models do not incorporate mutation of either hosts or microbes, for instance, nor are there temporal changes in processes whereby hosts acquire their microbes. Similarly, our models are based on discrete generations of hosts, and thus ignore the often very large differences in turnover rates of hosts and microbes. Nonetheless, these models, although simple, provide insights that are consistent with what we observe in nature, and also insights that are novel. In summary, our findings indicate that if the host provides resources to modulate or shape the microbiome, both its diversity and fitness decrease while the host’s fitness increases. Furthermore, under selective pressure, hosts are more likely to evolve towards providing resources to shape the microbiome when they exist in harsh environments with fewer beneficial microorganisms. Lastly, microorganisms not only need to adapt to the host’s internal environment but also contribute to the host in order to occupy a dominant position.

## Materials and Methods

The details of neutral and selection models have been described in Zeng et al. papers [8, 9].

### RPP

We have chosen to limit RPP to the TMS model by Zeng et al. [9]. This is because the HMS model by Zeng et al. already allows hosts to modify the fitness values of microbes through a heritable process. Although RPP does something similar, there are two major differences between HMS and RPP: (1) In HMS, hosts do not sacrifice resources, while in RPP, hosts sacrifice resources to modulate the microbiome; and (2) In HMS, the contribution to the fitness of OTUs varies within each host, whereas in RPP, the contribution to the fitness of OTUs varies based on the parameter *k*.

After incorporating RPP, the fitness values of OTUs in different hosts can thus be calculated as follows:

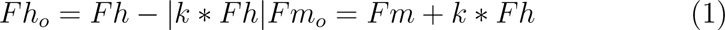

where *Fh_o_*represents the contribution to the host fitness of OTU in Host *o*, *Fh* represents the native contribution to the host fitness of OTU, *k* represents the *k* -value of Host *o*, *Fm_o_* represents the contribution to the microbial fitness of OTU in Host *o*, *Fm* represents the native contribution to the microbial fitness of OTU.

### Calculation of Microbial Diversity

In this study, *α*-diversity and *γ*-diversity (the overall microbial diversity from all hosts in the population) are calculated by scaled Shannon-Wiener index [35]:

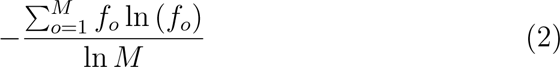

where *M* represents the total number of OTUs in the environment, *f_o_* represents the frequency of OTU *o* in microbial community.

And *β*-diversity (the change in microbial diversity between different hosts) is calculated by Bray-Curtis index [36]:

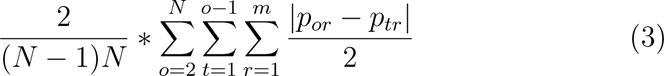

where *N* represents the number of hosts, *m* represents the total number of OTUs in all hosts, *p_or_* and *p_tr_* represent the frequency of OTU *r* in Host *o* and Host *t*.

### Model Implementation

In our RPP models, a random value, denoted as *k*, is assigned to each host from a uniform distribution ranging from 0 to 1 before the simulations. We adopted the same parameters as the selection model by Zeng et al. [9], including the number of hosts (*N* = 5000), the number of microbes carried by each host (*n* = 10^9^), the number of microbial OTUs (*M* = 150), the number of available traits (*j* = 25), and the number of traits carried by each microbial OTU (*i* = 5). The rationale behind these parameter settings has been explained in Zeng et al.’s paper [9]. However, we conducted simulations for fewer generations (*G* = 20000). We chose the number of generations based on the expectation under the Wright-Fisher model for haploid populations [37, 38] whereby the average time to the most recent common ancestor for neutrally evolving populations is twice the population size. Given that we have added selection in my model, we have determined that the equi-librium value of *k* is obtained before the 10,000th generation, and other indices stabilize before the 20,000th generation. This significantly reuces the computational time of each simulation. In addition, we used a grid approach to simulate all possible combinations of different selection intensities (*S_HS_* = 1, 5, 10, 100; *S_MS_* = 1.1, 5, 10, 100, 1000), different contribution rates (*x* = 0, 0.1, 0.2*, …,* 0.9, 1) from parent to offspring, and different contribution rates (*y* = 0, 0.1, 0.2*, …,* 0.9, 1) from pooled environment to mixed environment for simulation. Each combination was repeated 5 times (*R* = 5).In total, we performed 24200 simulations.

All figures were generated using the R programming language, and the simulations were performed using the Java programming language. The model was run on the New Zealand eScience Infrastructure (NeSI) high-performance computing facilities.

## Acknowledgements

We thank Qinglong Zeng for assistance with the original code, and Steve Wu and Anna Santure for helpful comments on an earlier draft of this work. Funds for this research were provided by The University of Auckland, New Zealand.

## Data Availability Statement

The code and dataset of this study are available at github.com/zjzxiaohei/Resource-Provisioning-Model.git.

