## Supplemental Figures for "Models of microbiome evolution incorporating host resource provisioning"

### Supplemental files 1: Figure S1

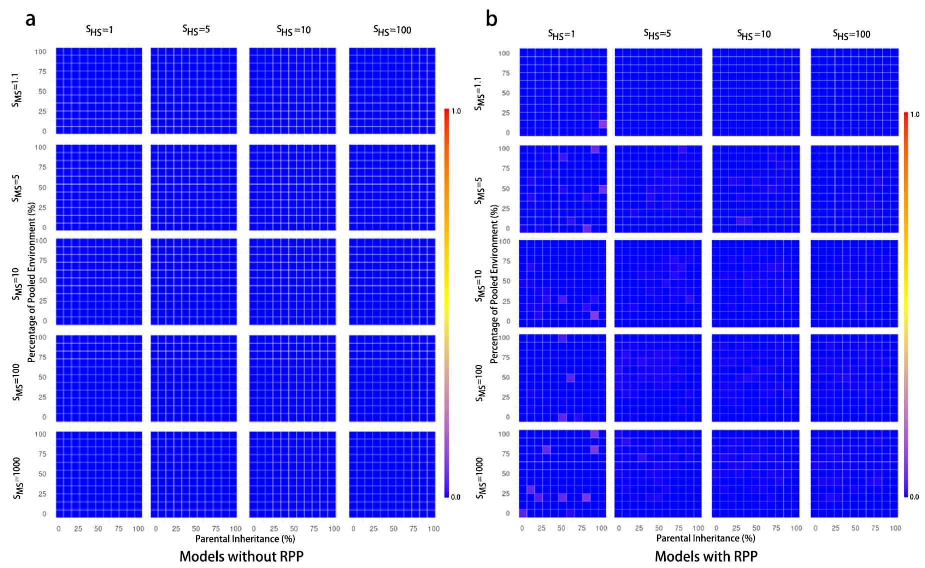


Figure S1: Pattern of $\beta$-diversity without and with RPP. Each heat map represents its corresponding combination of HS and MS. There are five levels of MS: 1.1, 5, 10, 100, 1000, and four levels of HS: 1, 5, 10, 100. For each heat map, the horizontal axis is the parental contribution to the host (i.e., parental inheritance, PI), and the vertical axis is the parental contribution to the environment (i.e., the percentage of pooled environment, PE). The scale of the horizontal and vertical axes of each heatmap is linear, from 0 to 100. The color bars on the right side represent the corresponding diversity values (blue for low diversity, yellow for medium diversity, and red for high diversity).

### Supplemental files 2: Figure S2

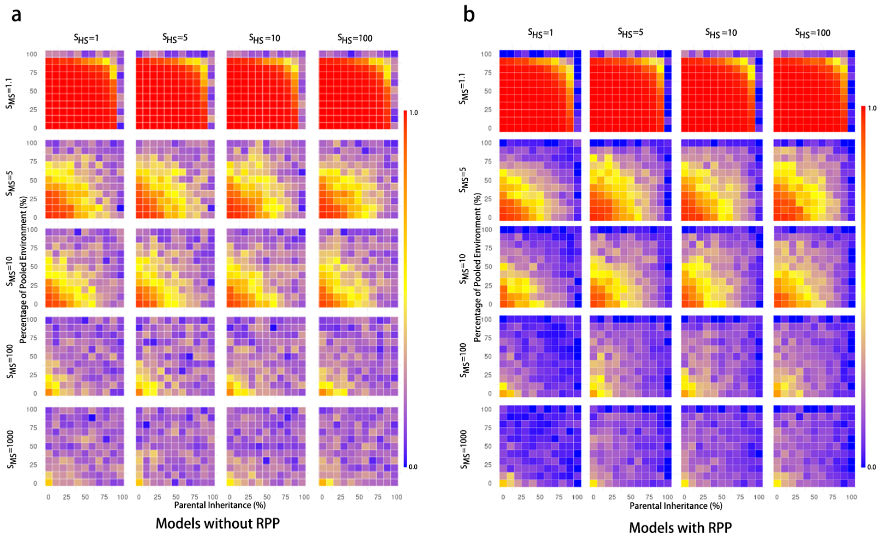


Figure S2: Pattern of $\gamma$-diversity without and with RPP. Each heat map represents its corresponding combination of HS and MS. There are five levels of MS: 1.1, 5, 10, 100, 1000, and four levels of HS: 1, 5, 10, 100. For each heat map, the horizontal axis is the parental contribution to the host (i.e., parental inheritance, PI), and the vertical axis is the parental contribution to the environment (i.e., the percentage of pooled environment, PE). The scale of the horizontal and vertical axes of each heatmap is linear, from 0 to 100. The color bars on the right side represent the corresponding diversity values (blue for low diversity, yellow for medium diversity, and red for high diversity).
